# Studying self-assembly of norovirus capsid by a combination of *in silico* methods

**DOI:** 10.1101/2024.01.21.575142

**Authors:** Jean-Charles Carvaillo, Thibault Tubiana, Sella Detchanamourtty, Stéphane Bressanelli, Fernando Luís Barroso da Silva, Yves Boulard

## Abstract

Understanding how macromolecular assembly occurs is a fundamental and challenging problem because spontaneous, precise assembly is at the center of most biological processes. It is an elaborate process that requires non-covalent stable interactions between partners to stabilize the desired architecture for a specific purpose. One of the advantages of virus models is that under adequate conditions capsid proteins can be efficiently assembled *in vitro* in the absence of any other component, providing simplified experimental models that can be rigorously characterized. The present study aims at describing the initial steps of molecular self-assembly of norovirus-like particles (NoVLPs, composed solely of the major norovirus capsid protein VP1), by combining *in silico* computational approaches to explore complementary physical properties. We show that this strategy allows not only recapitulating but also revising a former NoVLP assembly model. Our approach can be applied and extended to other problems in macromolecular assembly.

## Introduction

Large macromolecular complexes may consist of hundreds of individual components, including mainly proteins, nucleic acids, and lipids, to perform many vital tasks for the cell. Among the best known, we can cite the ribosome to synthesize proteins or spliceosome to catalyze pre-RNA splicing [1]. Although these macromolecular complexes’ final structures can be solved with X-ray crystallography or cryo-electron microscopy, their formation process is difficult to study and might be specific to physico- chemical conditions inconsistent with these experiments. Complex formation is an elaborate process that requires determining the order of assembly and investigating the biomolecular interactions among all partners. Moreover, it is a dynamic process and the interactions between binding molecules can serve to stabilize a preferred conformation to favor the binding of the next partner. To study those assembly pathways, virus capsids can be perfect study objects because they are large complexes made of multiple copies of a few proteins packaging the viral nucleic acid. In the simplest case, the capsid is made of a single type of protein that can self-assemble (homoassociation) even without the nucleic acid component [2].

Viruses are obligate intracellular parasites, meaning that viruses rely on intracellular resources and must hijack the cellular machinery of a host. They can invade and propagate into all types of life forms, including animals, plants, and bacteria [3]. Their replication is both a complex and straightforward mechanism. It is based on the implementation of their information into host cells. To do so, viruses are organized in two essential components: **(i)** The first corresponds to the information, i.e. the genetic material. It can be found as single or double-stranded nucleic acid (DNA or RNA) in one or several polynucleotide chain(s); **(ii)** The second is a protein shell that plays a major role in the protection and other biological functions such as transport and regulation of expression of this genetic information. A capsid containing the genetic material is formed from numerous copies of a single or a few protein sequences; A third component may be added, it corresponds to **(iii)** additional external layers encasing the capsid and that may include a membranous envelope acquired from a host cell membrane [4]. This study focuses on the understanding of the capsid assembly process and especially on the formation of assembly intermediates. Most capsids can be classified into two classes of geometry: rodlike or spherical [4]. Rodlike capsids usually adopt a helical symmetry that can accommodate in principle any length of nucleic acid. As for spherical capsids, they usually adopt icosahedral symmetry restrained by regular polyhedron properties. There are exactly 60 identical pieces in an icosahedron, but regular icosahedral capsids can actually include a multiple T of 60 protein subunits, thus allowing larger shells to be made with the same polypeptide length. For particular values of the triangulation number T (e.g. 1, 3, 4 …), the 60xT protein subunits can form the icosahedral shell by assuming T very similar (though not identical) conformations [5].

Both theoretical models and experimental techniques such as magic angle spinning nuclear magnetic resonance [6], time-resolved small-angle-X-ray scattering (TR-SAXS) [7,8], size exclusion chromatography [9], light scattering experiments [10] or native mass spectrometry [11–13] have been used to study assembly mechanisms of viral capsids. It should be noted that none of these experimental techniques allows the detection of all assembly intermediates. It remains difficult to parse all capsid assembly pathways and products that occur over large ranges of length and time scales (Ångström to micrometer and picosecond to hours), and most of which are transient [14]. To overcome this problem, several theoretical models at both nanometer spatial and nanosecond time scales have been developed. For example, mathematical models to study nucleocapsid assembly kinetics were established to determine the evolution of intermediates concentrations as a function of time [15]. Two major kinds of models, one to predict the assembly of empty capsids [hereafter called virus-like particles (VLPs)] and the second to describe the assembly around their nucleic acid (nucleocapsids) were developed. Both kinds of models are usually based on nucleation-growth schemes, where the limiting step is the formation of the initial nucleus, although cooperative models are also considered [4]. In nucleation-growth models, the nucleus corresponds to the smallest intermediate with a probability greater than 0.5 of growing to the complete capsid before disassembling [4,16,17]. Capsid assembly, particularly VLP assembly, can be also studied with particle-based dynamics simulations. Molecular dynamics simulations of capsid subunits can be computed to monitor their dynamics, thermodynamic features and interactions between them [18–20]. Monte Carlo methods can also be used for the last two tasks. Defining intermediates as particles or coarse-grained (CG) models enables faster sampling due to reduced degrees of freedom while maintaining the main features of the system. With these *in-silico* techniques, plausible assembly models have been defined [21,22], especially on T=3 VLP self-assembly [18]. Another example of *in-silico* CG capsid self-assembly studies is on human immunodeficiency virus 1 (HIV-1) VLP [20]. Numerical methods are high-performance computing techniques contributing to a better understanding of the assembly, biomolecular interactions and dynamics of highly multimeric structures and many other aspects of the viral life cycle [23]. The microsecond simulation of hepatitis B virus (HBV) T=4 VLP at an atomistic level can be cited as an example of a huge system performed on a supercomputer. The solvated and neutralized HBV VLP is composed of almost 6 million atoms [24]. This all-atom simulation indicates that even fully assembled capsid is capable of asymmetric distortion.

In 1999 Prasad and coworkers solved the Norwalk virus VLP structure at atomic resolution by X-ray crystallography (PDB id 1IHM, pH 4.8, resolution of 3.40Å) [25]. The structure showed that the Norovirus VLP is a classical T=3 icosahedron with 180 copies of the major protein VP1 in three quasi-equivalent sets A, B, and C, with the 60 ’A’ molecules around fivefold axes and the 60 ’B’ and 60 ’C’ molecules alternating around quasi-sixfold axes [5]. After analyzing the 3D structure, Prasad *et al.* proposed that **(i)** The Norovirus capsid assembly unit is a dimer of VP1; **(ii)** A pentamer of A-B dimers (POD) would be a major intermediate in the assembly of the 90-dimer Norovirus capsid. From this conclusion, an assembly model was suggested starting with the addition of 5 C-C dimers at the 5 positions around the POD, thus an isotropic growth from the POD nucleus. It is important to note that numerous experimental studies have reported that disassembly of several Norovirus VLPs (NoVLPs), including Norwalk virus VLP, indeed yields dimers of VP1 [12,26,27], for a review see ref. [13]. From these dimer solutions, T=3 VLPs can be reassembled *in vitro*, but for some Noroviruses smaller (presumably T=1) or much larger VLPs can also be assembled, *e.g.* [27]. The plasticity of capsid size of the Norovirus VP1 is also manifest in the fact that recombinant VLPs can display different T numbers from the normal T=3 of norovirus capsids, e.g. T=1 or T=4 [28]. This plasticity is likely due to the Norovirus VP1 architecture, with a flexible articulation between the N-terminal S (shell) domain, which makes up capsid-forming contacts between VP1 dimers, and the C-terminal P (protruding) domain, that provides the major VP1 dimerization interface.

Native mass spectrometry analyses of the Norwalk VLP in partial disassembly conditions showed oligomers in two size ranges, the smaller size oligomers ranging from 2 to 18 VP1 [11]. Subsequent ion mobility-mass spectrometry data were consistent with a five-dimer nucleus and further revealed that the smaller-range oligomers were consistent with extended sheet-like VLP subassemblies, and thus possible assembly intermediates [12]. Direct evidence for an assembly intermediate came from a TR-SAXS self-assembly study of a bovine norovirus VLP [29]. The whole millisecond to hours, nanometer-resolution kinetics were ascribable to only 3 species: **(i)** a form factor corresponding to the VP1 dimer, **(ii)** a form factor corresponding to a 10- or 11-dimer intermediate, and **(iii)** a form factor characteristic of the T=3 VLP. The form factor of the intermediate was interpretable as an elongated, stave-like subassembly and inconsistent with the isotropic assembly model proposed by Prasad *et al.* for the Norwalk VLP [25], but consistent with the extended sheet-like VLP subassemblies of Uetrecht *et al.* [12]. Thus, the experimental data points to an assembly pathway for the NoVLP where long-lived intermediates grow from a POD, but anisotropically rather than isotropically as proposed by Prasad *et al*.

To improve our comprehension of the formation of a stave-like intermediate from the VP1 POD, we developed an *in-silico* computational strategy, combining both all-atom and coarse-grained molecular dynamics, docking calculations, and electrostatic analysis. Our results show how the local symmetry breaks may foster anisotropic growth from an initial symmetric nucleus and suggest that the molecular model initially proposed to account for the stave-like intermediate needs to be revised. During this modeling process, we take advantage of the knowledge of correct solutions extracted from the capsid structure to assess the quality of intermediate results. We thus show how reliable the results can be in the absence of this *a priori* knowledge.

## Materials and methods

### Starting models

The completed asymmetric unit model of Norwalk Virus VP1 capsid protein was modeled based on crystallographic capsid structure (PDB id 1IHM, resolution of 3.40 Å, pH 4.8) [25]. The X-ray structure of chain A (or C) has been determined from residues 29-520 and chain B comprising residues 10-520 of the 530 residues. The N- and C-terminal missing residues (respectively 28 and 10 for chain A and 10 and 10 for chain B), which correspond to the flexible part of the molecule, were completed with Modeller 9.16 program [30]. The final models for the N- and C-termini were as predicted unstructured. After an energy minimization, they were carefully checked visually and with PROCHECK [31]. In the end, we obtained the all-atom complete models of both VP1 A-B and C-C dimers and the pentamer of A-B dimers (POD). This study was conducted before the emergence of the Alphafold program [32] but as we could not ignore its arrival during the writing of this manuscript, we also used it to predict the folding of both monomer and dimer models. The results confirmed that the missing residues of the crystal structure are unstructured, as they were predicted with low confidence scores (pLDDT values) in our final models.

### Assembling Study Method

The strategy used to study the formation of the complex molecular structure with computational methods is summarized in Figure 1. It has been applied to study the assembly of the Norwalk virus empty capsid (VLP). We can subdivide the cyclic process into four steps: **(1)** The all-atom starting model (POD) was converted to Martini 2.2 coarse-grained (CG) structure to accelerate structure sampling of the large complex. Partial charges were assigned assuming protonation states corresponding to the pH 7. This pH 7 CG structure was used to initiate at least 10 microseconds of production time of MD simulations with Gromacs (version 5.1.4 and 2016.4 [33]) after the proper equilibration; **(2)** After clustering the MD trajectory, a representative structure was selected, usually in the last cluster (see results); **(3)** This structure was then converted back to an atomistic model to allow docking of a new molecular brick (crystallographic VP1 dimer without the termini), and **(4)** the best docking solution was selected based on both the energy criterion and the root mean square difference (RMSD) in coordinates between a docking pose and the “exact” solution (assumed here to be the crystallographic structure obtained at pH 4.8). The corresponding POD+D (dimer) was then extracted from the crystal structure and termini were added before feeding into the next cycle. We also tried using directly the new structure issued from docking to initiate a new cycle but found this alternate strategy less suitable (see results).

**Figure 1.**
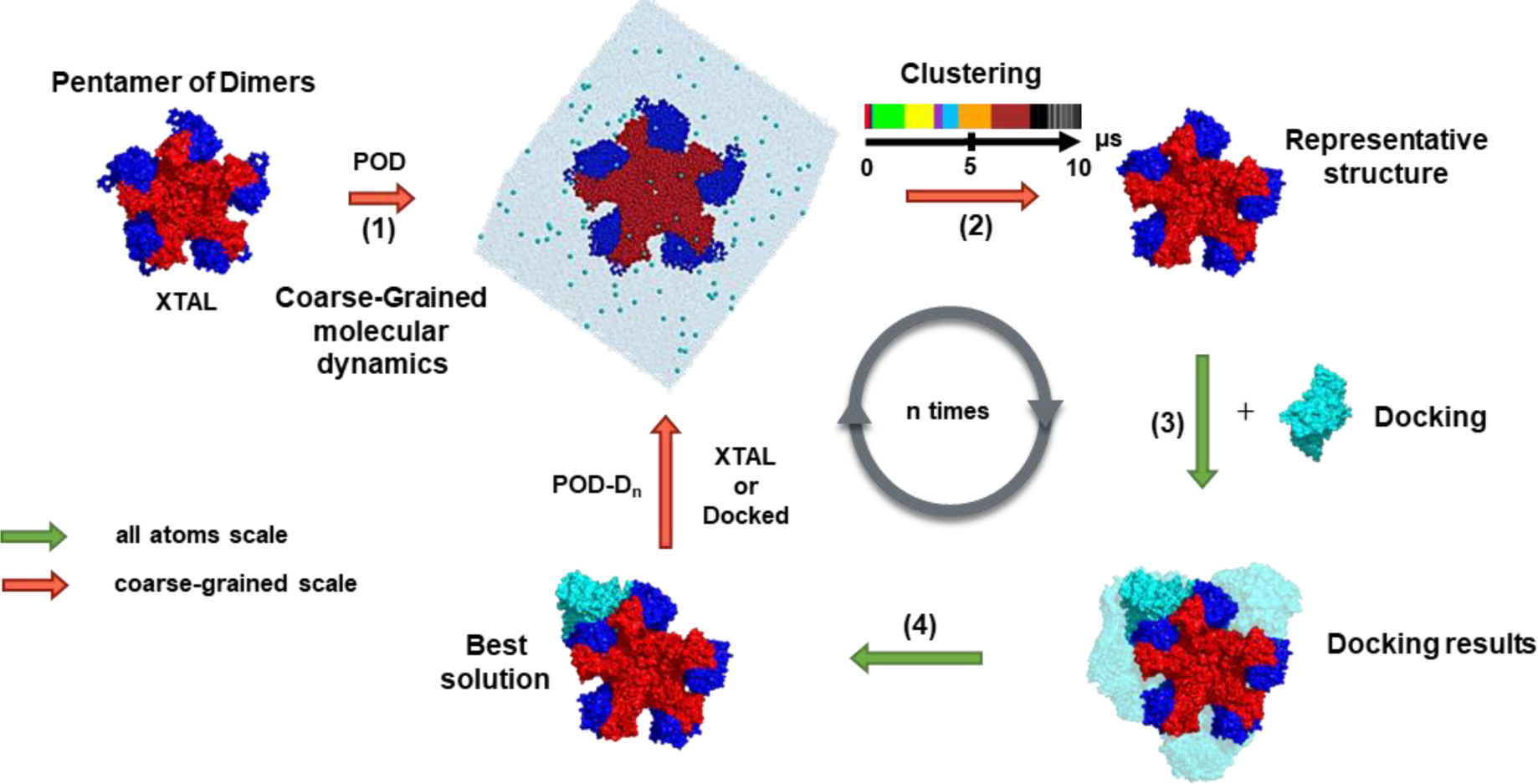
Computational strategy for studying the growth of the Norovirus capsid. The starting structure used is a pentamer of dimers (POD). For each dimer, chain A is colored in red and chain B in blue. Dimers added during the docking trial are colored in cyan. Our strategy has 4 steps: (1) the all-atom model is converted to a coarse-grained model, (2) a coarse-grained molecular dynamics simulation is performed, (3) docking of a new dimer is performed, (4) the best solution is retained to initiate a new cycle. The molecular dynamics simulation and clustering steps are at a coarse-grained scale symbolized by orange arrows. The docking step is performed at an all-atom scale (green arrows).

### Molecular dynamics calculations

We computed molecular dynamics (MD) simulations with the GROMACS simulation package version 5.1.4 and version 2016.4 [33] and the MARTINI 2.2 force- field [34,35] coupled to ElNeDyn elastic network to maintain secondary and tertiary structures [36]. To conserve independent flexibility and movement of S and P domains, we used the domELNEDIN algorithm to delete elastic bounds between these two domains [37]. A first steepest-descent energy minimization in vacuum on CG models was performed, Coulomb interactions within a cut-off radius of 0 to 1.2 nm and non-bonded Van der Waals (Lennard-Jones) interactions within a cut-off radius of 0.9 to 1.2 nm were used. Systems were neutralized with Na^+^ ions and solvated with MARTINI standard water combined to a low dielectric constant (ε = 15). Solvation was performed under periodic conditions with a non-cubic box with a distance between the protein and the edge of the box of at least 3 nm. The structures obtained after a second energy minimization were used to initiate MD simulations. All the MD simulations were performed at neutral pH and with the standard protocol in the NPT ensemble with temperature and pressure set to 300K and 1 bar, respectively. Charges were kept constant at their values assigned at the input phase. The covalent bonds were constrained using the LINCS algorithm [38]. Our protocol includes one equilibration step (2 ns) with the velocity rescale thermostat (τ_T_ = 1ps) [39] and Berendsen coupling algorithm (τ_P_= 4 ps) [40], followed by 2 relaxation steps (2ns each) with the Parinello-Rahman barostat (τ_P_ = 12 ps) [41]. During the equilibration and first relaxation step, coordinates were restrained with a force of 1000 kJ.mol^-1^ with a timestep of 10 fs. For the second relaxation step and production, restraints were removed and the timestep increased to 20 fs. VMD 1.9.3 1. [42] was used for RMSD calculations and Gromacs tool “rmsf” for the root mean square fluctuation (RMSF). To assess the robustness of our results, some CG simulations were replicated to confirm the observed trends. Additionally, we performed complementary simulations, for the POD system, with antifreeze particles in the Martini solvent with a proportion of 10% to prevent freezing effects in the CG water models.

### Clustering analysis

We used our *in-house* program TTClust [43] to cluster the trajectories with hierarchical clustering. The hierarchical distance is computed from the pairwise RMSD matrix Ward variance minimization algorithm [44]. The optimal number of clusters is estimated with the elbow method and a threshold of 0.995 with K-means clustering [45]. The representative frame is defined as the structure with the lowest RMSD against all the frames which belong to the same cluster.

### Docking procedure

Coarse-grained representative structures obtained after the clustering step are converted to all-atom structures with a going backward mapping dedicated tool [46] to allow docking procedure. Because backmapping is not a trivial task since it means reintroducing lost degrees of freedom, we checked the accuracy of the all-atom structures in various aspects including the atomic contacts at the protein-protein interfaces and the side chains construction. A protein-protein rigid-body docking (using fast Fourier transforms) was then performed on ClusPro 2.0 webserver with standard parameters [47] on a receptor (previous assembly iteration) and its ligand (VP1 dimer). After docking, ClusPro 2.0 computed several low-energy clusters that contain similar structures and extracted representative conformations. To discriminate docking solutions, we used the ClusPro 2.0 balanced scoring scheme and cluster centers PIPER energy functions [48]. In addition to the balanced score provided by ClusPro, we implemented a local RMSD metric using the crystallographic reference (PDB id 1IHM). The receptor used in the docking process (after MD simulation) is fitted to its parent crystallographic reference at each possible docking interface. To do so, we successively select the backbones of S domains (residues 30-220) lining each of these interfaces in the crystal structure. After superposition, we compute RMSD between residues 30-220 of the docked dimer and of the crystallographic dimer at this interface. After trying all possible interfaces of the current intermediate, the docked dimer is then assigned to the crystallographic dimer position with the smallest RMSD and this value is assigned as the local RMSD of the docked solution. We implemented this method as a tool based on PyMOL [49] functionalities. The method allowed us to test each crystallographic interface to identify which one is closest to the docked dimer and to sort and classify them. We plotted the local RMSDs on the y-axis and the corresponding docking interface on the x-axis. The number of members in a cluster is represented by the size of dots.

### Complementary analysis based on electrostatic features

It is well-known that electrostatic interactions play a pivotal role in viruses [50] Given the large size of these systems, we employ a simpler yet effective physicochemical framework, aimed at capturing the main physics of the problem to complete our analysis. When a charged protein (e.g. a ligand) approaches or even binds to another protein (e.g., a receptor), its electrical charge can induce changes in the ionizable properties of the titratable amino acids on the protein especially the ones at its nearby surface. Of course, due to the long-range nature of electrostatic interactions, a strong electrostatic coupling between titratable amino acids can be observed between groups of the intra and inter-chains residues, and even buried ionizable amino acids can be affected by the visits of the other charged molecule during an observation time. Such shifts in the protonation states provide valuable physico- chemical information and can offer a pragmatic approach to mapping all the titratable protein residues that are electrostatically affected by such visits during a simulation run [51]. By the comparison of the partial charges of a protein immersed in an aqueous electrolyte solution (apo state) with their values in the presence of its charged ligand in the same physico-chemical conditions (pH, ionic strength, protein concentration, and temperature), such charge shifts are directly estimated and can be used to identify the key amino acids that interact with the ligand (in the present case, the dimer). For a pragmatic use of this information to determine the specific surface of the receptor where the ligand has more often visited, a weak electrostatic coupling between the superficial and buried titratable group is the ideal scenario. As stronger is this coupling, a higher number of deeper and/or just neighbors’ amino acids are perturbed by the ligand coming nearer the receptor, and consequentially less effective is the theoretical prediction to map the preferential binding spots at the surface level. Conversely, it will indicate the main titratable groups responsible for the interactions of these two molecules. The partial charges of all macromolecular systems at pH 7 and 150 mM of 1:1 salt were calculated by the fast proton titration scheme (FPTS), a constant-pH (CpH) Monte Carlo (MC) simulation scheme for biomolecular systems [52]. FPTS was performed both for the apo state (POD in the absence of the ligand) and during the POD-dimer complexation (POD+D→POD-D) processes [53]. In the latter case, the separation distance and orientation of the molecules can affect the ionization process allowing the mapping of the involved titratable groups. We implemented a visualization and an analyzing PyMOL tool to exploit resulting partial charges (in elementary units) variations (δq) that are explained further in the results section. Free energies of interactions for the dimer-dimer associations as a function of their separation distances (r) were also calculated at different salt concentrations (I) and pH conditions using the CpH MC FORTE approach [54]. Besides the dimer structure given by the PDB id 1IHM (GI), additional MC runs were performed with the PDB id 6OTF (resolution 3.1Å, pH 5.75) to allow some additional comparisons with another human norovirus from a different genogroup (GII). Some key physico-chemical properties such as the total net charge numbers (Z), the total charge regulation capacity (C), and the total dipole moment numbers (µ) were computed as well by the FPTS.

The structure of the Norwalk virus T=3 VLP (49% sequence identity with the bovine Norovirus VP1, 59% in the S domain) was used as it was the only available atomic structure at the time we started this study (PDB id 1IHM). We extracted a POD as the most likely assembly nucleus from the available experimental data [12,25]. The three-dimensional structure was then carefully completed and minimized, and protonation states were assigned at pH 7. This choice was based on the fact that despite experimental data on VLP being acquired at pH 6 for the kinetics study [29] or pH 9 for the ion mobility-mass spectrometry study [12], bovine Norovirus and Norwalk virus capsids both self-assemble and are fairly stable at pH 7 [11,26,27]. Moreover, as for all single-stranded, positive-sense RNA viruses, Noroviruses assemble in a cytosolic compartment that is presumably close to neutral pH. The all-atom model was converted to Martini 2.2 CG structure and relaxed. The relaxed structure was used to initiate MD simulations (20 microseconds of production time) with Gromacs. Trajectory clustering indicates the system evolves directionally, visiting successive conformations that are substantially different from the initial symmetric POD and stabilizing several microseconds in each without coming back to a previous conformation (see Figure 2). Examination of a representative frame of the two final conformations after 10 µs and 20 µs of simulation shows that in both cases the initial symmetry of the POD is broken. In the symmetric POD extracted from the NoVLP, the five outlying S domains from molecules ’B’ are regularly spaced, leaving five identical spaces between two successive domains that in the NoVLP are occupied by five C-C dimers. In contrast, after MD simulations two successive outlying S domains are either considerably nearer to each other or farther away. This behavior is found consistently in the replicas for the POD simulations. This prompted us to consider these particular, transiently stable conformations as the likely targets of further capsid growth, The representative CG structures after 10 µs or 20 µs were then converted to all-atom structures and relaxed to allow docking of a new dimer [46]. The docking approach was also performed on the symmetric POD from the NoVLP to assess the consequences of this consistent symmetry break. The 30 representative structures output by ClusPro were then visually analyzed. Some of them are satisfactory, the new dimer docks between two outlying S domains of the POD as expected but some of them are not compatible with a new intermediate to form the final capsid (some examples are shown in Figure S2). To discriminate between the acceptable solutions and the false ones, we developed an analyzing tool. It is based on the calculation of a local RMSD between expected positions according to the X-ray structure and the docking solution considered (see methods for details). The cut-off for this local RMSD was determined primarily visually on the graph. However, as depicted in Figure 2B, there is a well-defined gap between good and bad RMSD values. An RMSD of 16 Å certainly permits us to validate good solutions (threshold represented by a dashed line) and it excludes bad or false positive energetic solutions. This strategy was further validated by the fact that we observed a very good correlation between the best energy structures and the lower RMSD docking results (see Figure 2C). The final results are shown in Figure 2 for the POD structure. Clearly, for the symmetric POD extracted from the crystal structure, the five favorable positions (between two successive outlying S domains) for docking a new dimer are equivalent (see Figure 2A) whereas for the asymmetric POD resulting from 10 μs of the MD simulations, one potential docking position is never found and the 4 others are not equivalent (see Figure 2B). Positions 1 and 2 seem to be the best. After 20 μs of the MD simulations (see Figure 2D), the situation was slightly different since 2 docking positions appeared favorable, positions 2 and 4. At this stage, we retained position 2 to add a new dimer at the POD and to continue the study. Starting from this new structure named POD-D, we used the same strategy as described above to add a new dimer. The results are shown in Figure 3.

**Figure 2.**
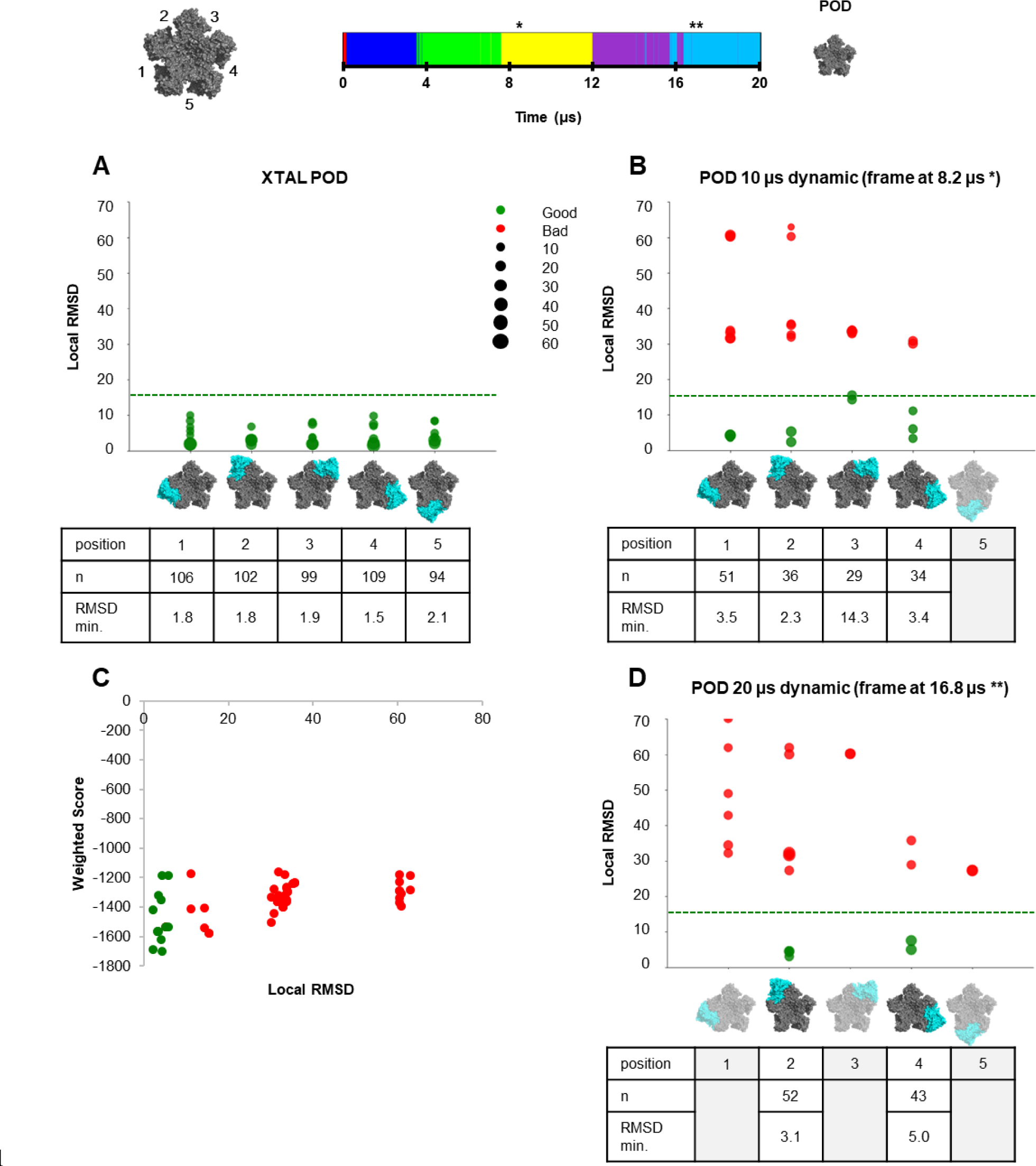
Docking solutions onto POD structure. Top panel, the timeline of the simulation. The asterisks denote the representative frames of two transiently stable conformations in a 10 µs coarse- grained molecular dynamics simulation (*) or in the same simulation extended to 20 µs (**). **(A)** Docking results onto the POD structure extracted from the X-ray crystal (**B**) Docking onto the representative structure after 8.2 µs. (**D**) Docking onto the representative structure after 16.8 µs. Each docking solution output by ClusPro was checked against each of the five possible positions represented below the plots, both visually and by computing a local RMSD (see materials and methods). The solution was then assigned to the position with the minimum local RMSD. The horizontal dashed line at 16 Å indicates a cutoff beyond which the docking solution does not match an acceptable solution, while it always does below this cutoff (bad solutions are in red circles and good solutions are in green circles). The sizes of circles are proportional to the number of ClusPro members for each docking solution. The tables report the total numbers of good solutions for each position 1 to 5. (**C**) PIPER weighted score computed by ClusPro as a function of local RMSD.

**Figure 3.**
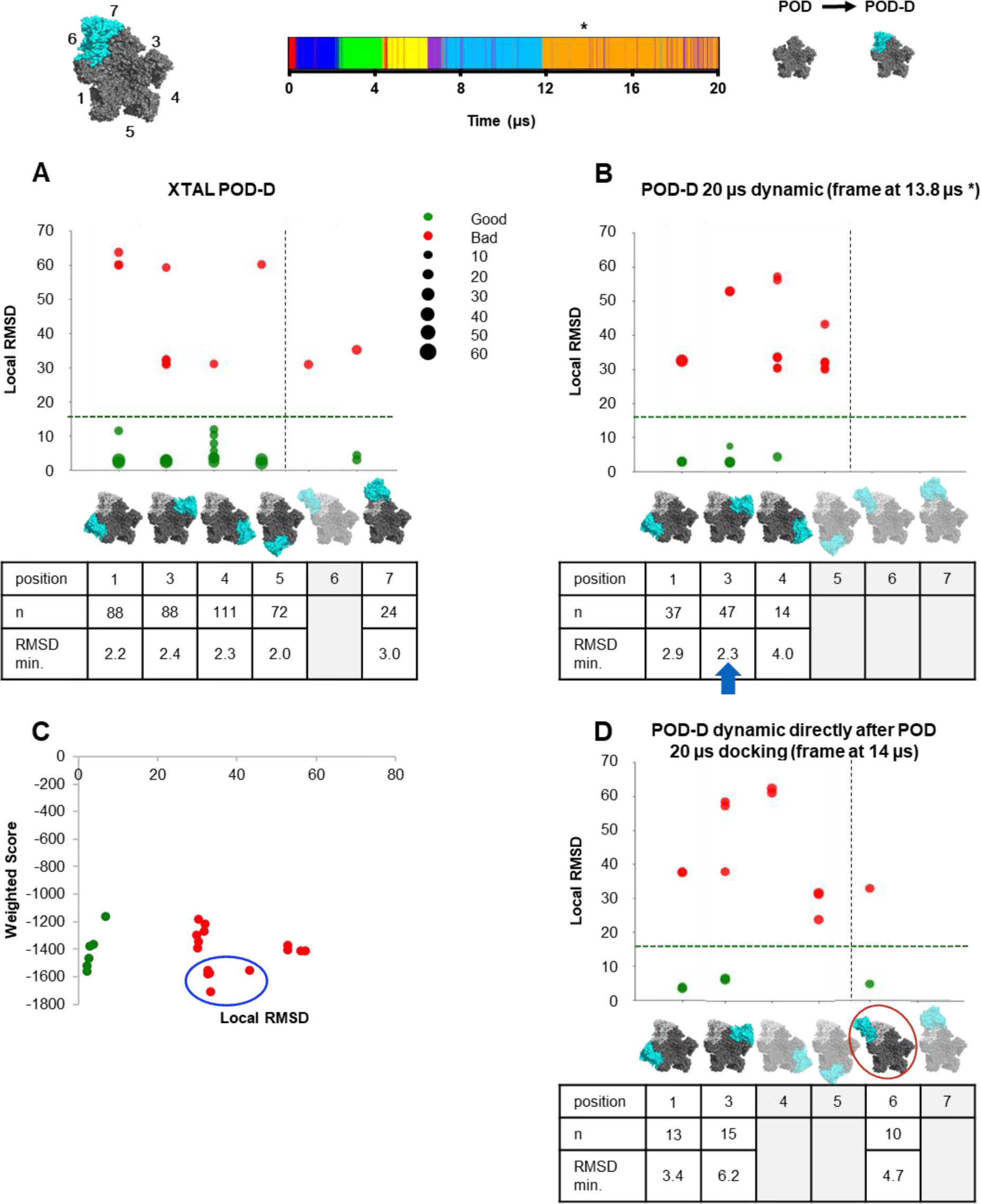
Docking solutions onto POD-D structure. Vertical lines separate isotropic growth from anisotropic growth. **(A)** Docking results onto the POD-D structure extracted from the X-ray crystal (**B**) Docking results onto the representative structure after 13.8 µs coarse-grained simulation. The best solution (blue arrow) will be used to study isotropic growth. (**C**) PIPER weighted score computed by ClusPro as a function of local RMSD. (**D**) Docking results onto the best previous docking solution selected after a 10 µs simulation (POD docking solution at position 2 - Fig 2B) which was used to initiate a second 10 µs molecular dynamics. The docking solution at position 5 (surrounded by a red circle) which corresponds to an anisotropic growth was selected to explore anisotropic path.

Based on the POD-D structure, we expected six favorable positions to dock a new dimer. Four of them correspond to the free remaining positions of the initial POD structure. The other two possibilities correspond to docking on the supplementary dimer. As shown in the graph given in Figure 3A, only five of them are identified as acceptable solutions if we consider the crystal capsid structure as the reference. After 20 μs of MD simulation, the number of favorable docking solutions for a new dimer is reduced to three (see Figure 3B). They correspond to free positions on the POD structure, the two positions adjacent to the first docked dimer and one opposite the first docked dimer. We observe that there is still a good correlation between the best energy structures and the lower RMSD docking results (see Figure 3C). However, some very favorable solutions for the energy criterion are now clearly excluded if we consider the RMSD criterion (blue circle in Figure 3C). At this stage, we retained the solution for which there were the most positive results with the lowest minimum RMSD (blue arrow Figure 3B). It corresponds to the structure in which the new dimer docks near the previous one and we name it POD-D_2_ 2-3 (structure POD with 2 docked dimers at positions 2 and 3). The results of the addition of a new dimer to this structure are shown in Figure 4. Without surprise, if we consider the X-ray structure, the three remaining positions of the initial POD structure are very satisfactory (positions 1, 4, and 5 in Figure 4A) resulting in an isotropic growth of the capsid. In contrast, two out of the four possible positions for anisotropic growth are favorable (positions 7 and 9 in Figure 4A) but with less positive solutions (47 vs 273). After the MD run, the results are significantly different (see Figure 4B) since only two positions are advantageous to dock a new dimer, the first corresponding to an isotropic growth (position 4 in Figure 4B) and the second to an anisotropic growth (position 8 in Figure 4B). If these solutions are numerically and energetically almost equivalent, the RMSD value is better for docking on the initial POD structure. As the structure grows, evidently, the best solutions do not correspond to the ones selected by ClusPro with only an energy criterion as shown in Figure 4C.

**Figure 4.**
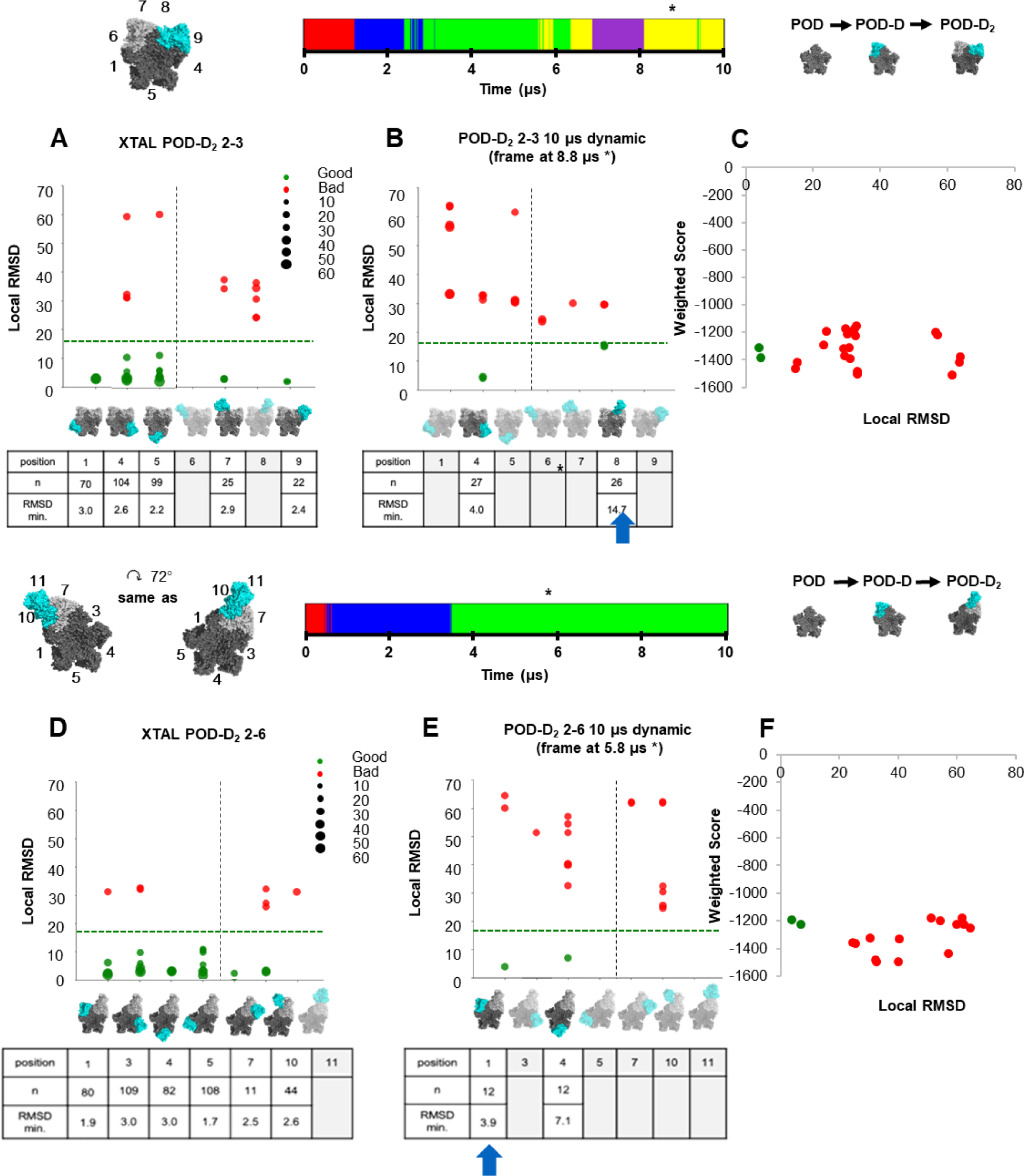
Docking solutions onto POD-D_2_ 2-3 structure. **(A)** Docking results onto the POD-D_2_ 2-3 structure extracted from the X-ray crystal or **(B)** onto the representative structure after 8.8 µs simulation. Vertical lines separate isotropic from anisotropic docking solutions. The green arrow represents a common docked structure between adjacent and continued POD-D_2_. (**C**) PIPER weighted score computed by ClusPro as a function of local RMSD.

As explained in the Materials and Methods section, we also tested an alternate strategy of keeping the best-docked solution to initiate a new cycle of docking an additional VP1 dimer, thus keeping the conformational changes introduced by the simulations. Compared to the approach described above, several drawbacks appear. Firstly, the curvature of the viral capsid is not maintained and secondly, defects accumulate, as the molecular assembly grows compared to the reference crystal structure. This accumulation of errors makes the selection of good docking solutions more and more difficult. However, at the start of the procedure, it is quite satisfactory. The docked solutions are interesting and close to those described above. In particular, at the POD-D step, a second pathway leading up to an anisotropic growth appeared (position 6 in Figure 3D). This solution, named POD-D_2_ 2-6, was observed with our first approach as shown in Figure 3A, but the docked dimer’s position was too far from the crystal structure to be retained. In general, isotropic docking solutions are preferred as shown in Table 1. But it clearly depends on our choice for the first docking dimer for which many good solutions are possible.

**Table 1:**
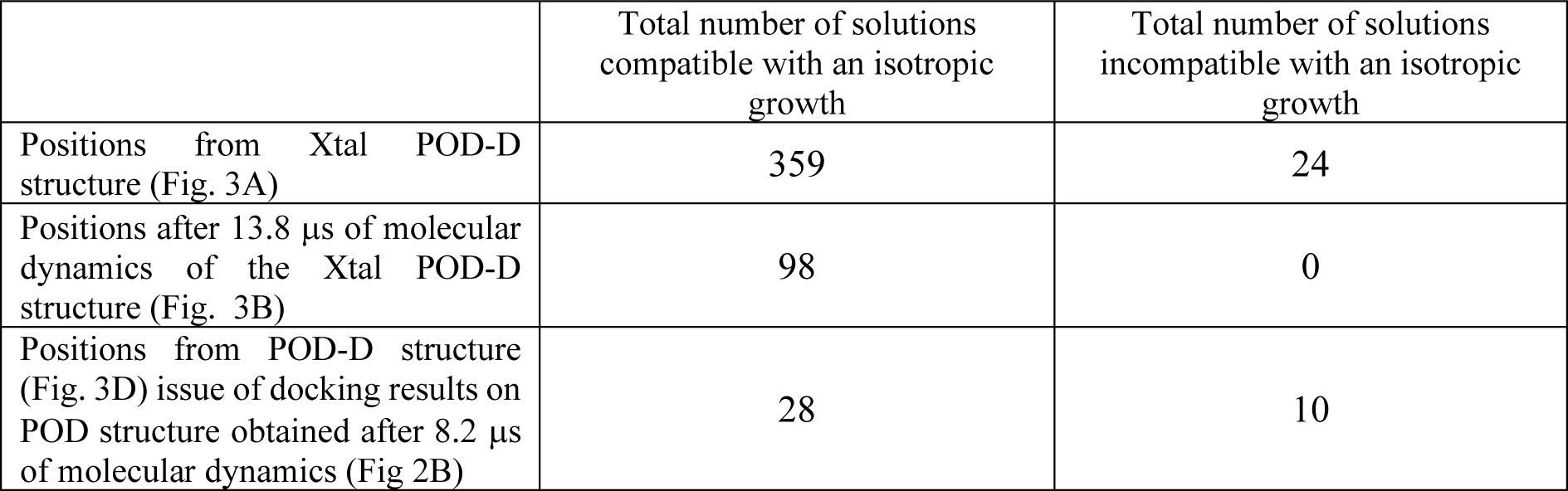
Comparison of POD-D docking results.

For the rest of the study, we explored the two paths as shown in Figures 4 and 5 for POD-D_2_ and POD- D3 steps respectively. The results are shown in Figure 6. All the potential docking solutions that remained to complete the initial POD structure are observed for the POD-D_2_ crystal structure as shown in Figure 4. It corresponds to positions 1, 4, and 5 in Figure 4A and positions 1, 3, 4, and 5 in Figure 4D. The solutions 1, 4, and 5 in Figure 4A are compatible with an isotropic growth for the assembly. The remaining solutions (positions 7 and 9 in Figure 4A and solutions in Figure 4D) are compatible with anisotropic growth. After the MD run, we observed only 4 favorable docking positions for the supplementary dimer; 3 of them correspond to a docking on the initial POD structure (position 2 in Figure 4B and 1 and 3 in Figure 4E) and only one to a docking on a new interface (position 6 in Figure 4B). These solutions are close for both RMSD and energy criteria as shown in Figures 4C and 4F. Interestingly, the two paths have a common POD-D_3_ solution, (position 6 Figure 4B and position 1 Figure 4E). The results concerning the next step (POD-D_3_ step) are shown in Figure 5 for both paths. If we continue the isotropic path from the crystallographic structure or after MD runs, a common solution appears (Figures 5A and 5B) which corresponds to the docking of a dimer at position 5 on the initial POD. It is even the unique good solution after MD run whereas 3 other docking emplacements are possible on the crystallographic structure (positions 1, 8, and 12 Figure 5A). From the anisotropic growth path, the number of good docking emplacements increases to 6, including 3 on the initial POD structure (positions 1, 4, and 5 in Figure 5D). After the MD runs, only 2 structures are selected, with an additional dimer at positions 4 and 5 on the initial POD (see Figure 5E). Interestingly, a common good docking solution appears at the POD-D_3_ step, whatever path we consider, it is the position 5 on the initial POD. At this stage, the best solutions selected on the basis of the local RMSD criteria do not necessarily correspond, as observed previously, to the solutions with the best energy (see Figures 5C and 5F). Now that we have explored the two paths (one compatible with an isotropic growth, the second with an anisotropic growth) to the POD-D_4_ intermediate, the question arises of the intermediate identified by TR-SAXS knowing that it is composed of 10-11 dimers. Our results show that it is possible to build a single POD-D_5_, combining the complexes obtained for the two paths and in agreement with the form factor determined experimentally. This intermediate is shown in Figure 6.

**Figure 5.**
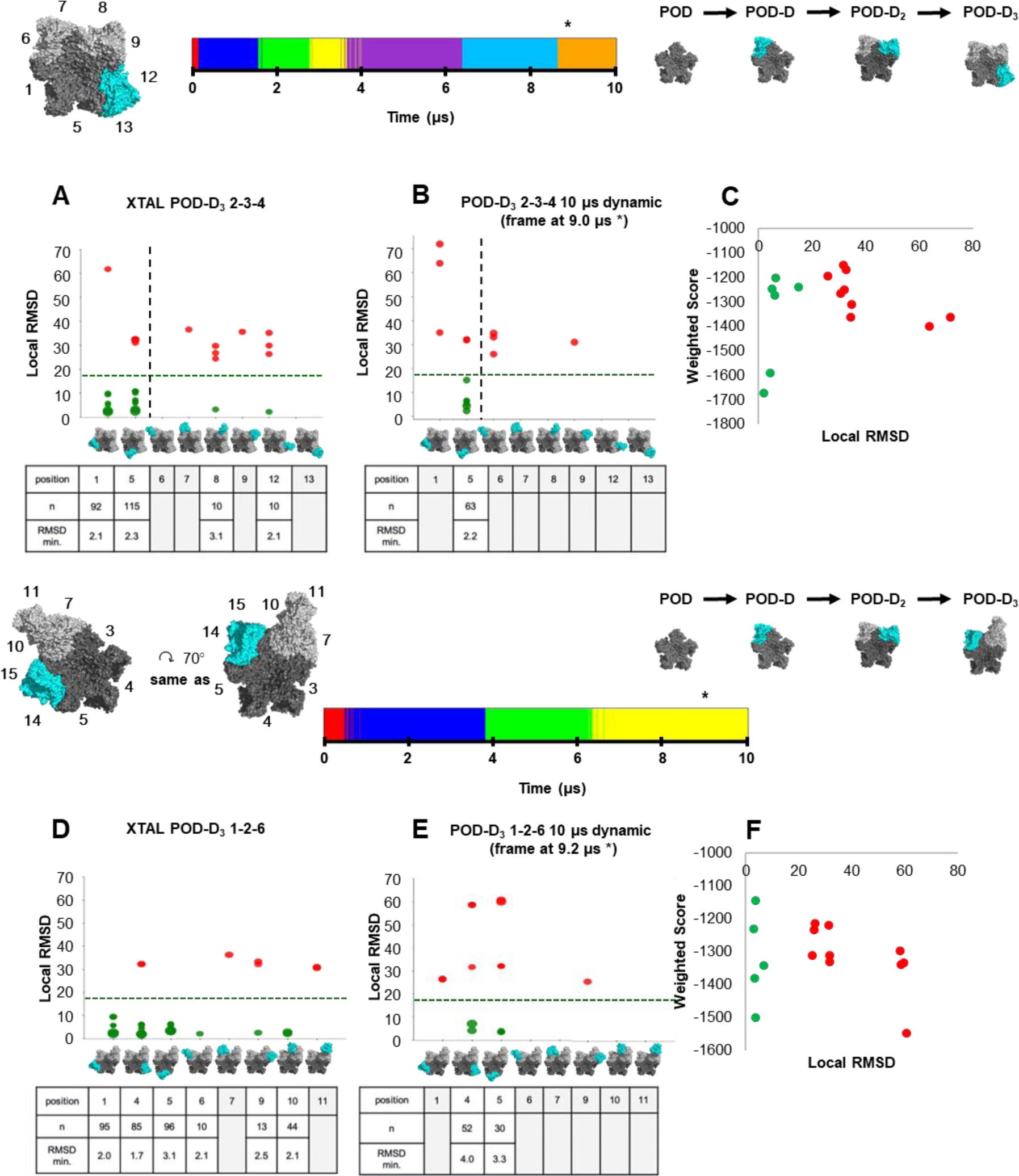
Docking solutions onto POD-D_2_ 2-6 structure. (A) Docking results onto the POD-D_2_ 2-6 structure extracted from the X-ray crystal or **(B)** onto the representative structure after 5.8 µs simulation. Vertical lines separate isotropic from anisotropic docking solutions. The green arrow represents a common docked structure between adjacent and continued POD-D_2_. (**C**) PIPER weighted score computed by ClusPro as a function of local RMSD.

**Figure 6.**
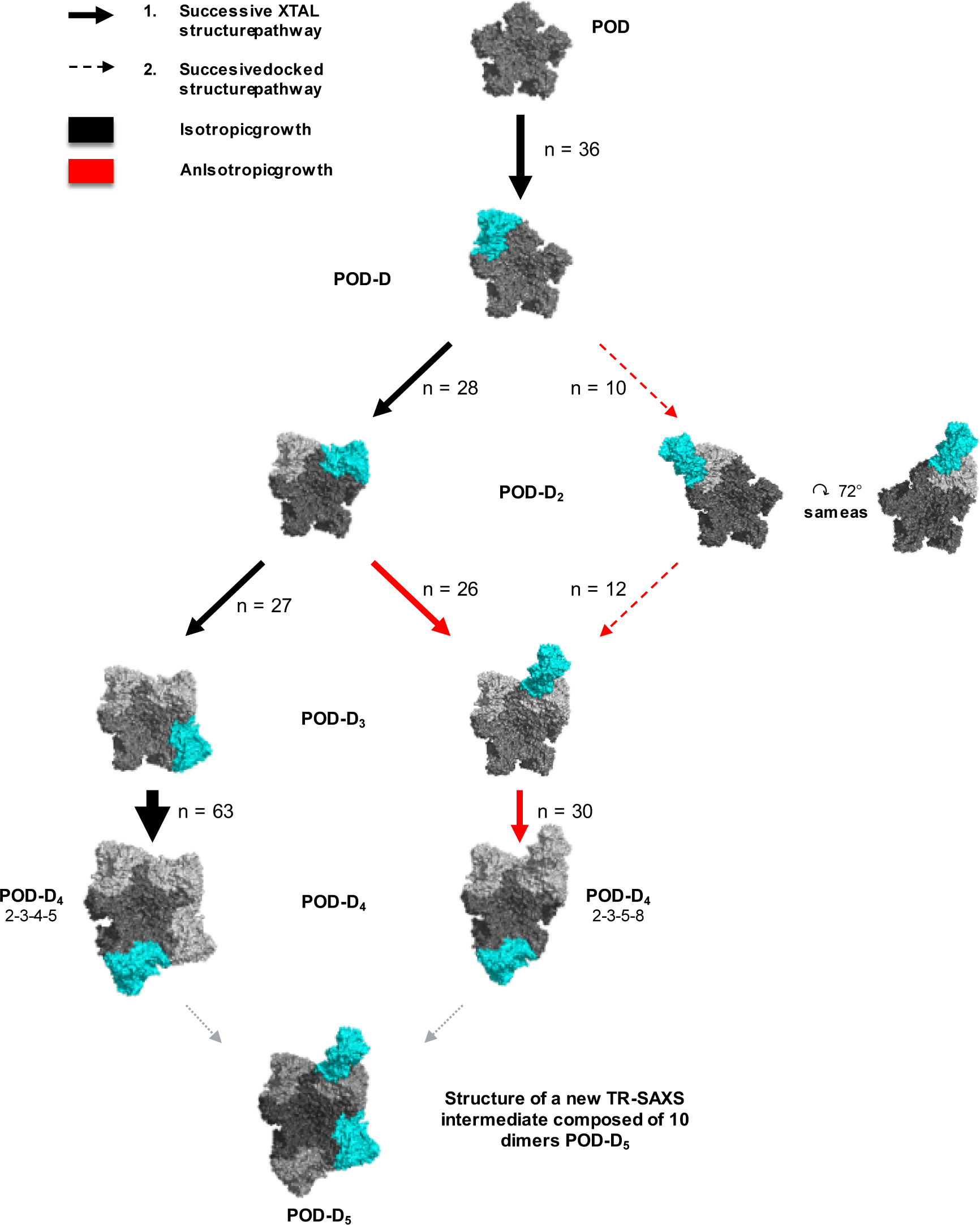
Growth pathways from the POD structure result from our docking strategy. The number n corresponds to the number of solutions at each step. By extrapolation, POD-D_5_ corresponds to a model compatible with the elongated assembly intermediate identified by Tresset *et al.* during the norovirus capsid self-assembly study [29].

The above procedure, summarized in Figure 1, thus allows several insights into how building of the Norovirus capsid by accretion of subunits may favor an initial anisotropic growth, by initial symmetry break and by convergence of several distinct pathways towards the same intermediate. However, it requires several simplifications that do not take into account some important parameters of Norovirus capsid assembly. One is the known fact that titratable groups of VP1 are important, as witness the modulation of Norovirus capsid assembly and stability by pH (Tresset et al., *Arch. Biochem. Biophys.* 2013). Another related issue is that these modulations are dependent on the Norovirus strain, with marked differences even in genogroups that both infect humans such as GI and GII[55]. To investigate the electrostatic features of VP1 self-interactions, as mentioned above, we used a computing solution following the strategy developed by the Barroso da Silva team, the PROCEEDpKa method [56]. The core idea of this approach is the identification of all titratable groups of the receptor that were perturbed by the visits and presence of the ligand. When the electrostatic coupling of these ionizable groups is not so strong, the *pKa* shifts can be used to identify the key amino acids responsible for a ligand-receptor association [56]. For the sake of comparison with the other simulation approaches and due to the larger size of the macromolecules involved in the present study, the PROCEEDpKa method was simplified here to analyze only the charges shifts at solution pH 7.0 instead of the *pKa* shifts. We used this method to determine if there was a correlation between the best docking results obtained using a constant-charge methodology (i.e. when the charges are assigned at the input phase for a given pH condition and do not change during the simulation run) and the electrostatic properties of these structures. We thus computed the shifts in charges of ionizable residues in the POD in the presence of a sixth dimer during a complexation study. This was compared with the charges obtained from a titration study in the absence of this sixth dimer (“apo” case). We first plotted these charges shifts near expected interfaces, with a cutoff chosen to exclude residues in nearby interfaces (see supplementary Figure S3). Whatever the system we studied (POD, POD-D, or POD-D_2_), we found no apparent differences between expected interfaces with this criterion (see supplementary Figures S4 and S5). On one side, this implies that the electrostatic coupling between the ionizable sites is quite strong for the POD system. It also indicates that both inner and superficial titratable groups can effectively participate in the interplay of the molecular interactions among the dimers. Indeed, the importance of electrostatics is well established experimentally for norovirus capsid dimers. For instance, the above-mentioned TR-SAXS study was performed with dimers dissociated at pH 9 and low salinity by a jump to pH 6 and salinity 55-105 mM [29]. We verified this phenomenon with the CpH MC [52,56] simulation method recently named FORTE (**F**ast cOarse-grained pRotein-proTein modEl) [54], and found that dimer-dimer interactions are indeed computed as repulsive at pH 9 and ionic strength (I) equals to 10mM and as attractive at pH 6 and I = 55mM (see the black and red curves, respectively, in Figure 7). On another side, it hampers the practical application of this approach to probe and reveal the existence of any preferential binding spots at the surface level for this particular system, if any.

**Figure 7.**
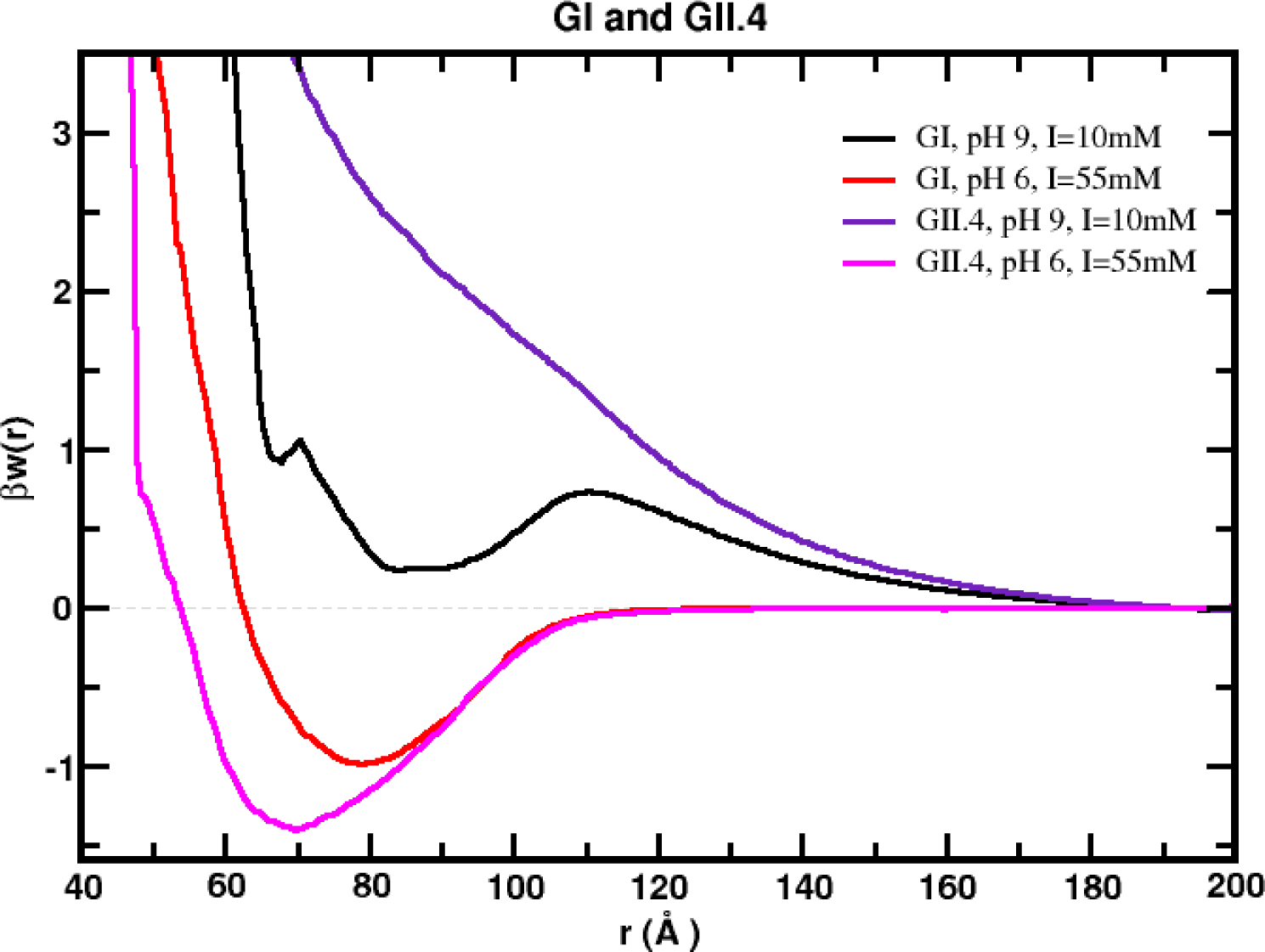
Free energy profiles for the dimer-dimer homoassociation. The simulated free energy of interactions [βw(r)] between the centers of mass of the two macromolecules at different physico- chemical conditions were obtained from CpH MC simulations [48,50]. These simulations started with the two macromolecules placed at random orientation and separation distance. The temperature was fixed at 298K. Free energies are given in k_B_T units (k_B_ is the Boltzmann constant). Data for the interaction of both the GI and the GII dimer homoassociation at pH 9 and I = 10mM and pH 6 and I = 55mM.

We next applied the FORTE approach to another norovirus VP1 dimer-dimer interaction, this one of genogroup II [28]. It is structurally almost identical to the GI dimer (RMSD of 0.9 Å between the two dimeric three-dimensional structures), but of high divergence at the sequence level (Identity of 42.2%). Furthermore, it is the closest in sequence (71.4%) to the GII Kawasaki strain, whose VLP was shown experimentally to be much more stable at high pH than GI strains [55]. The dimer-dimer complexation for GII is qualitatively similar to what was computed for GI (see Figure 7). Yet, it tends to be more attractive suggesting the formati.on of more stable capsids. This can be seen in Figure 7 for the comparison of the free energy of interactions for the dimer-dimer association [βw(r)] for both systems at pH 9 and I = 55mM. This result is consistent with the observation that some GII VLPs are more resistant to dissociation by high pH than other genogroups’ VLPs [13].

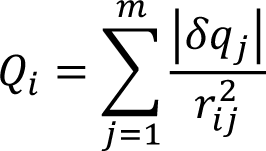

where *r_ij_* is the distance between *i* ^th^ dimer’s center of mass and the *j* ^th^ titratable residue’s center of mass. The green spheres correspond to correctly docked solutions. Orange and red spheres correspond to incorrectly docked solutions.

At this stage, we sought to combine the calculated shifts in charges with the docking results differently. As shown in Figure 8A for the POD system, we mapped on the 3D structure the charged shifted amino acids and the center of mass of all representative docking positions for the new dimer calculated with ClusPro. VP1 dimer is composed of two structural domains, the N-terminal shell domain (S) making up the protein capsid and the C-terminal protruding domain (P) making up the exterior surface of the shell (supplementary Figure S6), If one draws a plane separating domains S from domains P (Figure 8A, bottom), it is apparent that the most shifted POD residues (darkest grays) are generally on the side of P domains, while all ClusPro docking solutions are on the side of S domains, be they correct (green) or incorrect (orange and red) solutions. Reasoning that electrostatic interactions are long-range, and that there might be a strong coupling between titratable groups as mentioned above, we calculated, for each docking position *i*, a total electrostatic contribution of the POD residues Q_i_ based simply on Coulomb’s law (i.e. proportional to the magnitude of each shift and inversely proportional to the distance squared). Interestingly, we observed on average that the docking solutions corresponding to the correct addition of the next dimer (in green in Figure 8B) had higher Q_i_ than near misses (orange). However, the highest Q_i_ were computed for some particular docking positions in red Figure 8. Indeed, these last solutions correspond to positions where the dimer is docked in the center of the POD structure, near the 5-fold axis, which implies that the dimer is roughly at an equal distance from each of the five copies of any given residue in the POD. Consequently, its contribution to the charge shift of this residue is equivalent to each dimer belonging to the POD structure. Definitely, the good docking positions (in green) have always a high Q_i_ value.

**Figure 8.**
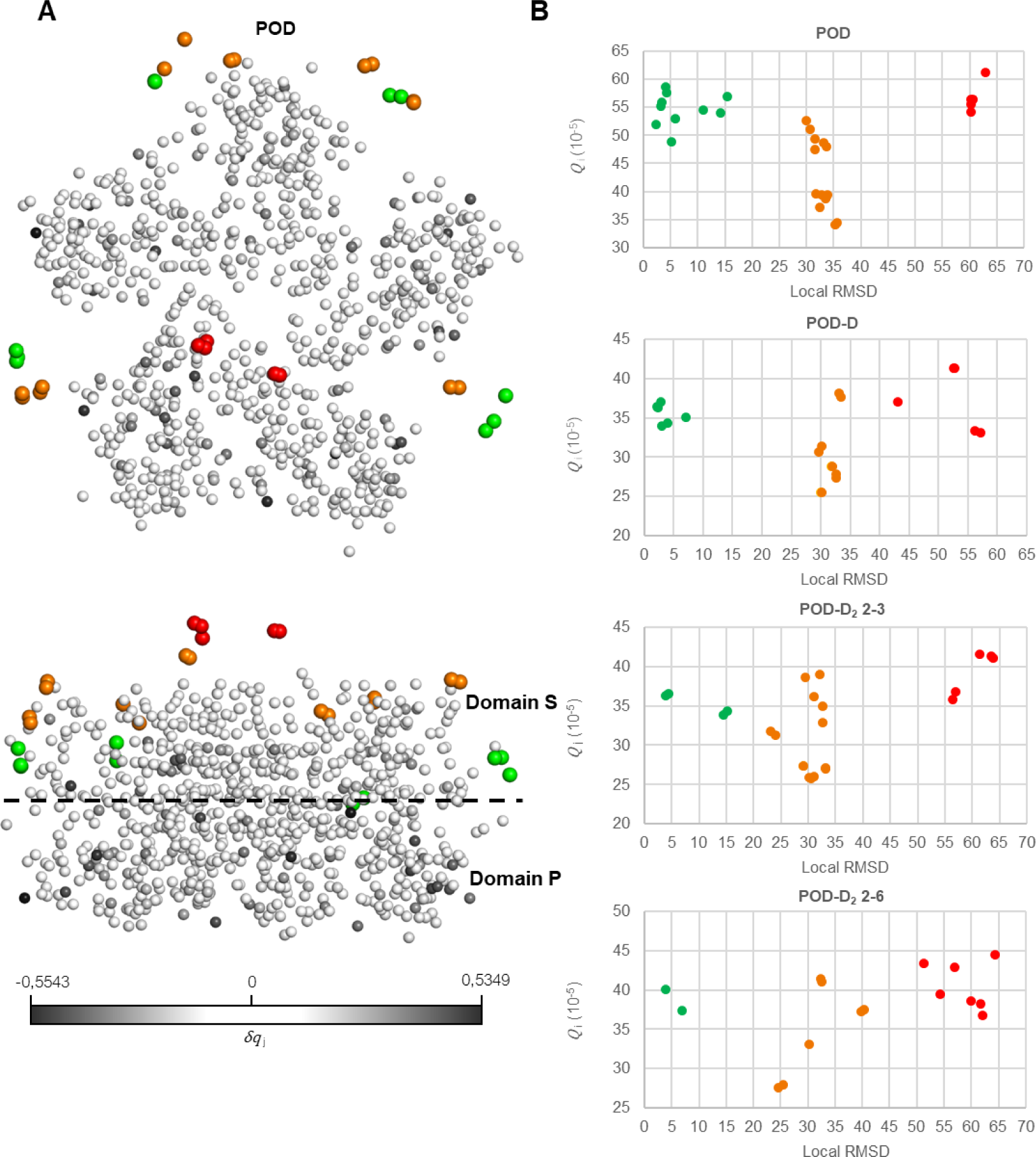
Relationship between docking solutions and titratable residues. (A) Localization of center of mass of docking solutions and titratable residues. (B) Total electrostatic contribution as a function of RMSD. For *i=1* … *n* docked dimers, and *j=1 … m* titratable residues, each with a shift *δq_j_*, we computed:

Thus, this combined analysis noticeably discriminates between correct and near-correct solutions at least under the assumptions behind this work. However, although the electrostatic analysis can identify the key amino acids for the association, the studied structures’ electrostatic variations are not sufficient to distinguish correct solutions from widely incorrect but geometrically central solutions for this system with such a strong electrostatic coupling among its ionizable groups. It demonstrates that the docking results we determined with our strategy (based on a constant-charge approach) could not be unambiguously reproduced with the sole mapping of the electrostatics features. This is also in accordance with the predominantly hydrophobic interfaces between domains S. Now, when we plotted the Q_i_ value versus the PIPER energy of ClusPro (Figure 9), we observed that the best energy docking positions have globally a high Qi value. It is particularly interesting because it suggests that without knowledge of the final solution (crystal structure), it may be possible to discriminate docking solutions based on the energy values combined with the electrostatic properties of the molecules. This result is very encouraging and demonstrates that it must be possible to describe the assembly mechanisms of biological molecular complexes by combining different simulation methods.

**Figure 9.**
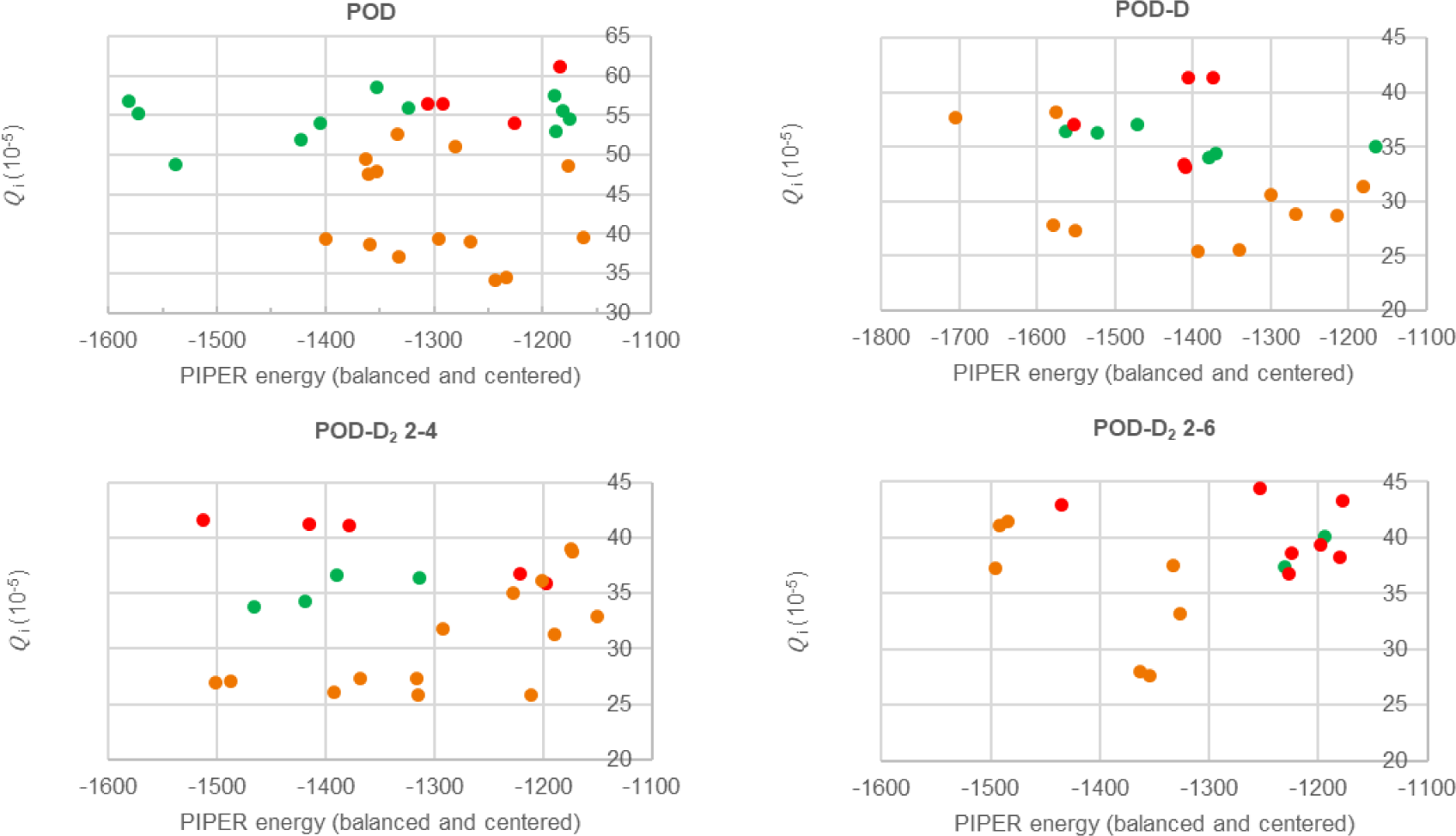
PIPER weighted score computed by ClusPro as a function of total electrostatic contribution for POD, POD-D, and POD-D_2_ systems. The green spheres correspond to correctly docked solutions. Orange and red spheres correspond to incorrectly docked solutions.

## Conclusion

Deciphering the assembly of macromolecular complexes remains a challenging question, but our work demonstrates that taking into account the dynamical properties of the molecules, both conformational changes, and variations in physico-chemical properties induced by the interactions between two molecules, are essential to select the best solutions proposed by docking. Indeed, based on *in silico* predictions combining molecular dynamics simulations, docking, and electrostatic properties, our results demonstrate that starting from the POD structure of the Norwalk VLP, an elongated intermediate similar to the one experimentally identified in the self-assembly kinetics of the bovine norovirus VLP [29] is accessible with our strategy. However, the model we are led to is not consistent with the one that was proposed in the former work: As an isotropic growth around the POD is favored in the first steps, our results rule out two PODs connected by a single C-C dimer. We propose two possible explanations for this discrepancy. The first possibility is that our simplified procedure does not capture some key property of bovine Norovirus VLP self-assembly. Apart from charge variation, the most obvious one is reversibility of subunit addition, an important feature of capsid self-assembly [57,58] that we do not consider when only adding VP1 dimers. But an alternate possibility is that the model of the intermediate proposed previously is not correct: So, pushing our reasoning, based on our new results, we propose that elongated intermediates composed of 10 dimers (Figure 6) may lock together to form a complete capsid. This hypothesis is very attractive because it allows a complete 90-dimer VLP to form with 9 intermediates of 10 dimers, without the need for *ad hoc* additional dimers. Furthermore, this solution remains compatible with previous experimental data on Norovirus VLP (dis)assembly. First, it is compatible with the form factor of the intermediate extracted from the TR-SAXS data (Figure 6). But it is also noteworthy that ion mobility-mass spectrometry analysis of partially disassembled Norwalk VLP found no species between a 9-dimer extended sheet-like oligomer and 20-, 30- and 40-dimer more capsid-like (spherical) objects [12], suggesting that 10-dimer elongated VP1 assemblies may indeed spontaneously assemble into partial icosahedral VLP. Furthermore, the plasticity of norovirus capsid size, seen for instance in the ability to form T=4 capsids (120 dimers) or smaller than 90 dimer assemblies [28], is better accounted for by such 10-dimer intermediates. In actual Norovirus assembly, the single viral genomic RNA, covalently linked to the VPg protein, is recruited to the assembling capsid most likely through the involvement of the basic minor capsid protein VP2. The known interaction of VP1 with VP2 could serve to trigger, accelerate and/or complete the assembly of nine 10-dimer VP1 subassemblies, thus combining capsid assembly with genome packaging in the absence of a direct interaction between the major capsid protein and the genome.

Our approach, applied here on the Norwalk virus VLP, is general and can be adapted to other problems of macromolecular assembly. Its most immediate use would be on other virus capsids relying on the self-assembly of one or a few proteins. From a biological point of view, capsid assembly and maturation correspond to the late stages of the viral life cycle which are essential for the formation of infectious viral particles. Consequently, the development of drugs that target viral capsid assembly is attractive. For instance, ’capsid assembly modulators’ that derail the self-assembly of the hepatitis B virus nucleocapsid by binding at dimer-dimer interfaces have been developed, which are now in clinical trials for the treatment of this major public health problem [59]. Our approach could help in understanding the molecular mechanisms underlying these interesting and important phenomena.

## Acknowledgments

This work was granted access to the high-performance computing resources of TGCC under allocation A0050710605, A0070710605, and A0090710605 made by GENCI. We also thank the CEA-CCRT infrastructure for giving us access to the supercomputer COBALT. J-CC was supported by a predoctoral fellowship from ANRS (France Recherche Nord & Sud SIDA-HIV Hépatites: FRENSH) and TT by the ANRS postdoctoral fellowship ECTZ189696, AAP 2018-1. F.L.B.S. is also deeply thankful to the support given by “Fundação de Amparo à Pesquisa do Estado de São Paulo” (Fapesp 2020/07158-2), the Conselho Nacional de Desenvolvimento Científico e Tecnológico (CNPq 305393/2020-0) and the Swedish National Infrastructure for Computing (SNIC 2022/5-601). This research was also done using computational services provided by the OSG Consortium, which is supported by the National Science Foundation awards #2030508 and #1836650.

## Author Contributions

**Conceptualization:** SB, YB

**Data curation:** JCC, SD

**Formal analysis:** JCC, SB, FLBS, YB **Funding acquisition:** SB, YB **Investigation:** JCC, TT, SB, FLBS, YB **Methodology:** FLBS, YB

Project administration: **SB, YB**

**Writing – original draft:** JCC, YB

**Writing – review & editing:** JCC, TT, SB, FLBS, YB

