## supplementary Figure for "Studying self-assembly of norovirus capsid by a combination of *in silico* methods"

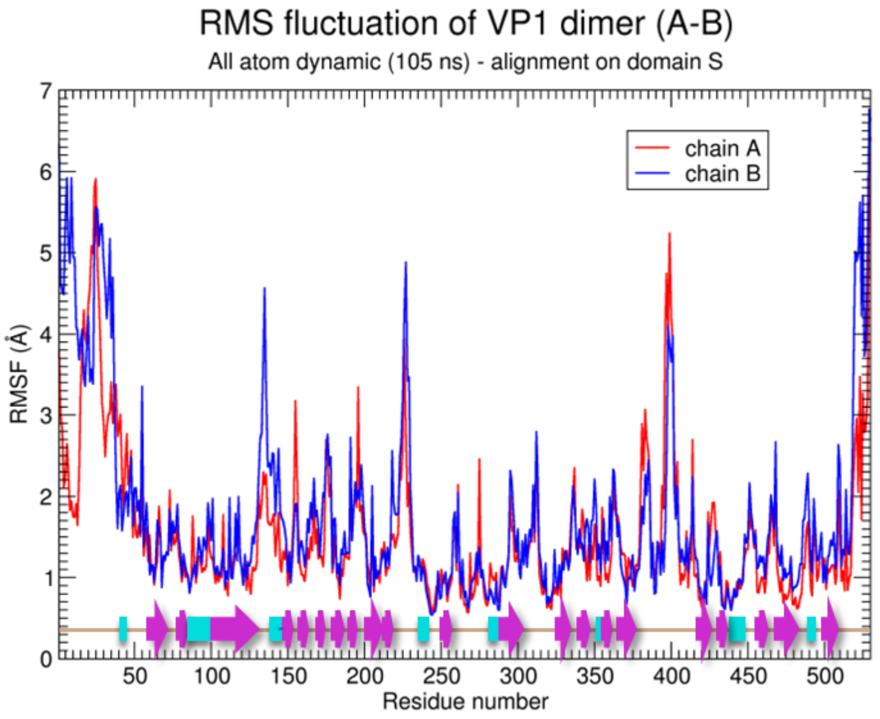


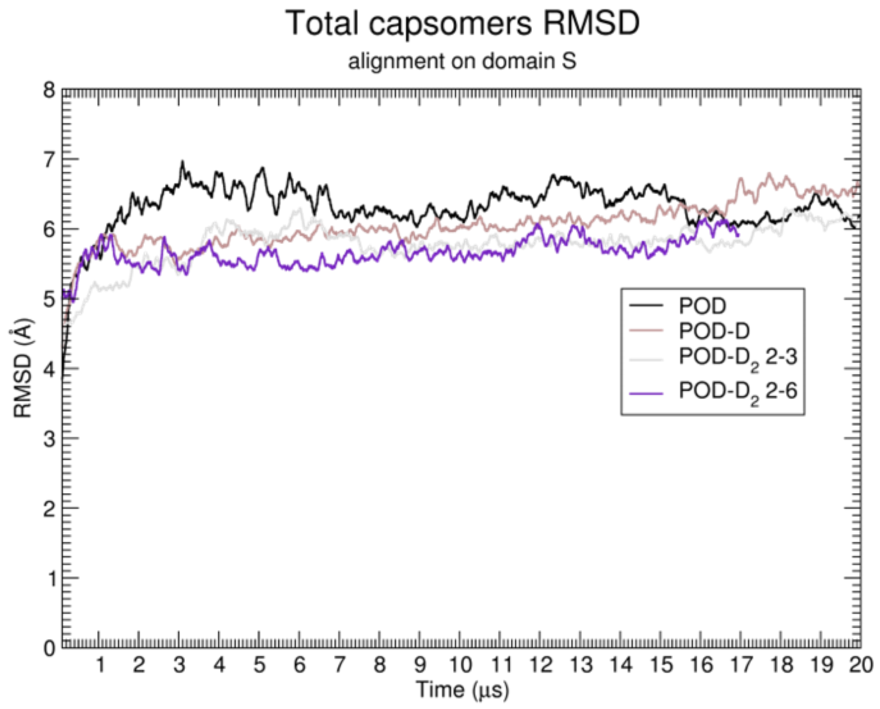


**Supplementary Figure 1.**

**Root Mean Square Flutuation and Root Mean Square Deviation.** RMSF values during 105 ns of the all atoms dimer structure and RMSD of coarse grained structures.

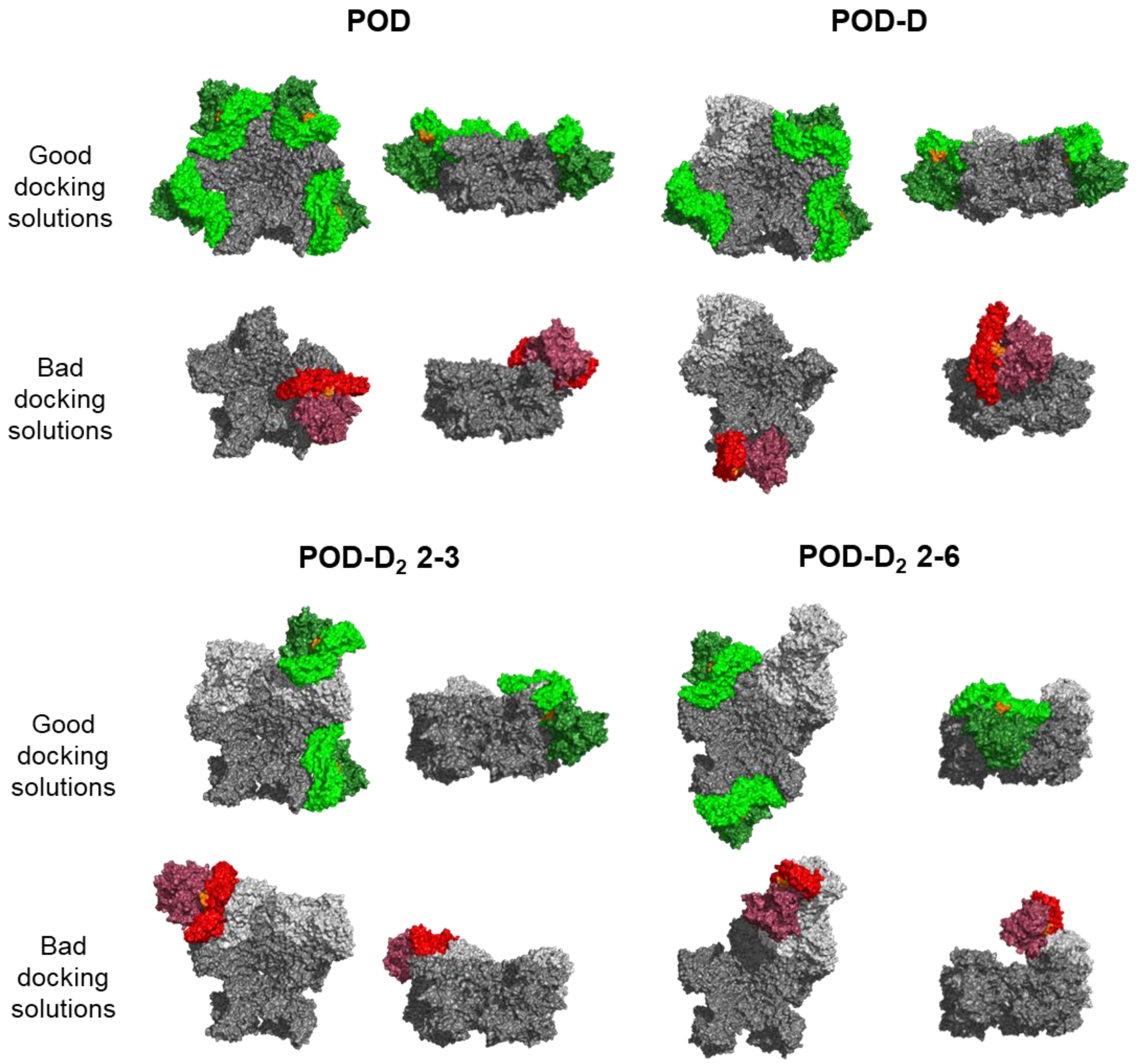


**Supplementary Figure 2.**

**Examples of good and bad docking solutions.** POD, POD-D POD-$D_2$ 2-3, and POD-$D_2$ 2-6 are colored in grey (POD in dark grey and additional dimers in light grey). Good docking solutions are shown in green and bad docking solutions in red (domains S are in light green or light red and domains P are in dark green or dark red).

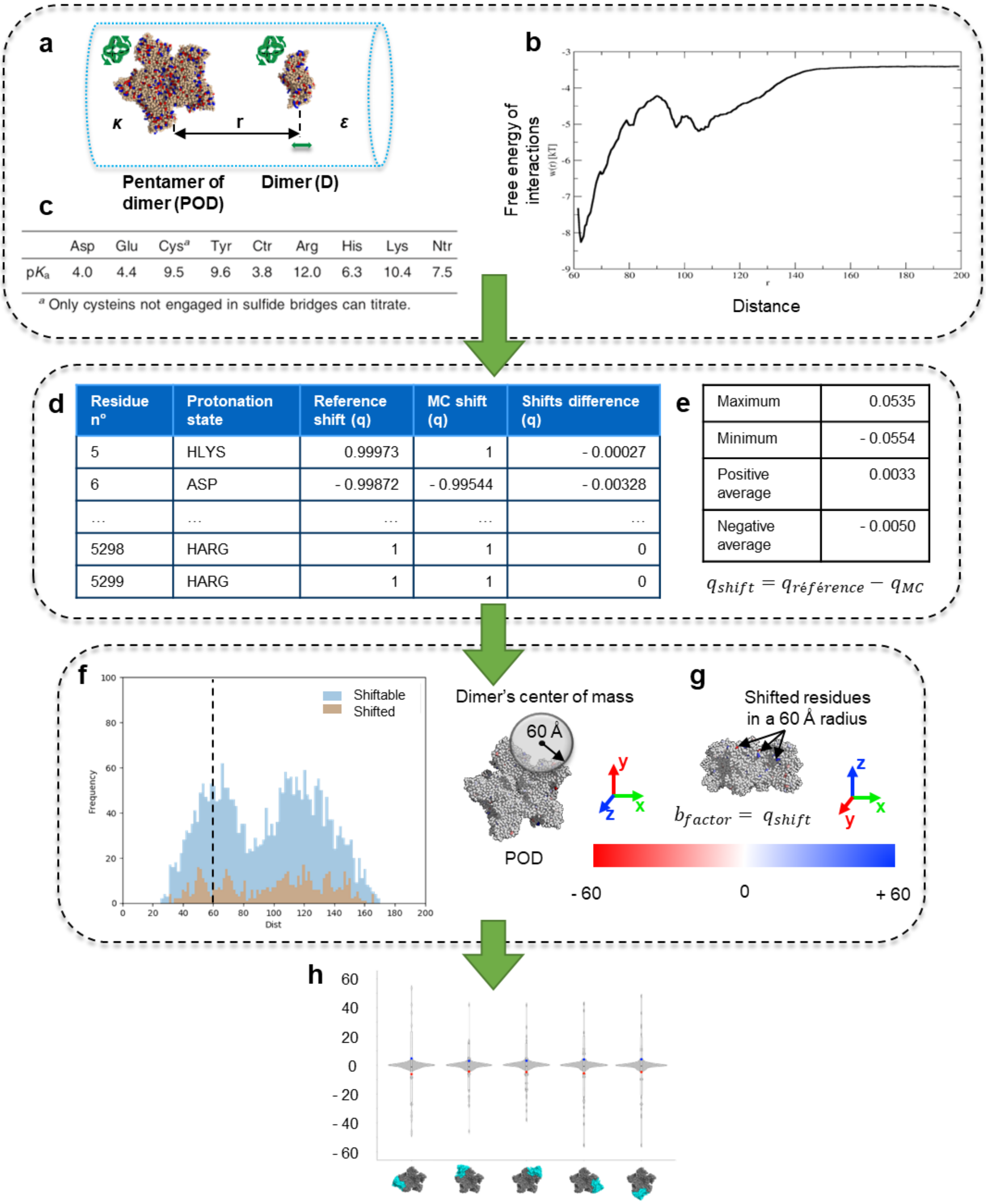


**Supplementary Figure 3.**

**Impact of dimer approach on ionizable residues of the receptor**. Charge shifts were calculated on POD, POD-D, POD-$D_2$ systems as illustrated in (**a**, **b,** and **c**). Charges are firstly computed on the system without ligand (column "Reference shift (q)" and secondly in presence of a new dimer (column "MC shift (q)" in (**d**)). It allows calculating the shift of ionizable residues (column "Shifts difference (q)" in (**d**)).

$$q_{shift} = q_{reference} - q_{MC}$$

To identify significant shifts, we decided to use positive and negative averages as threshold (**e**). $q_{shift}$ upper positive average and below negative average are considered significant shifts. To discriminate and quantify charge shifting at a specific interface of docking positions, we calculated both distributions of the number of ionizable residues and of significantly shifted residues as a function of the distance from the center of mass of the concerned dimers in the X-ray structure of the capsid (**f**). It allows us to select a radius of 60 Å around a docked dimer's center of mass as a good limit to define the docking interface (**g**). Finally, violin plots (it combines boxplot and density plot) were used to visualize our data (**h**). cf. **Suppl. Fig 4 and 5**.

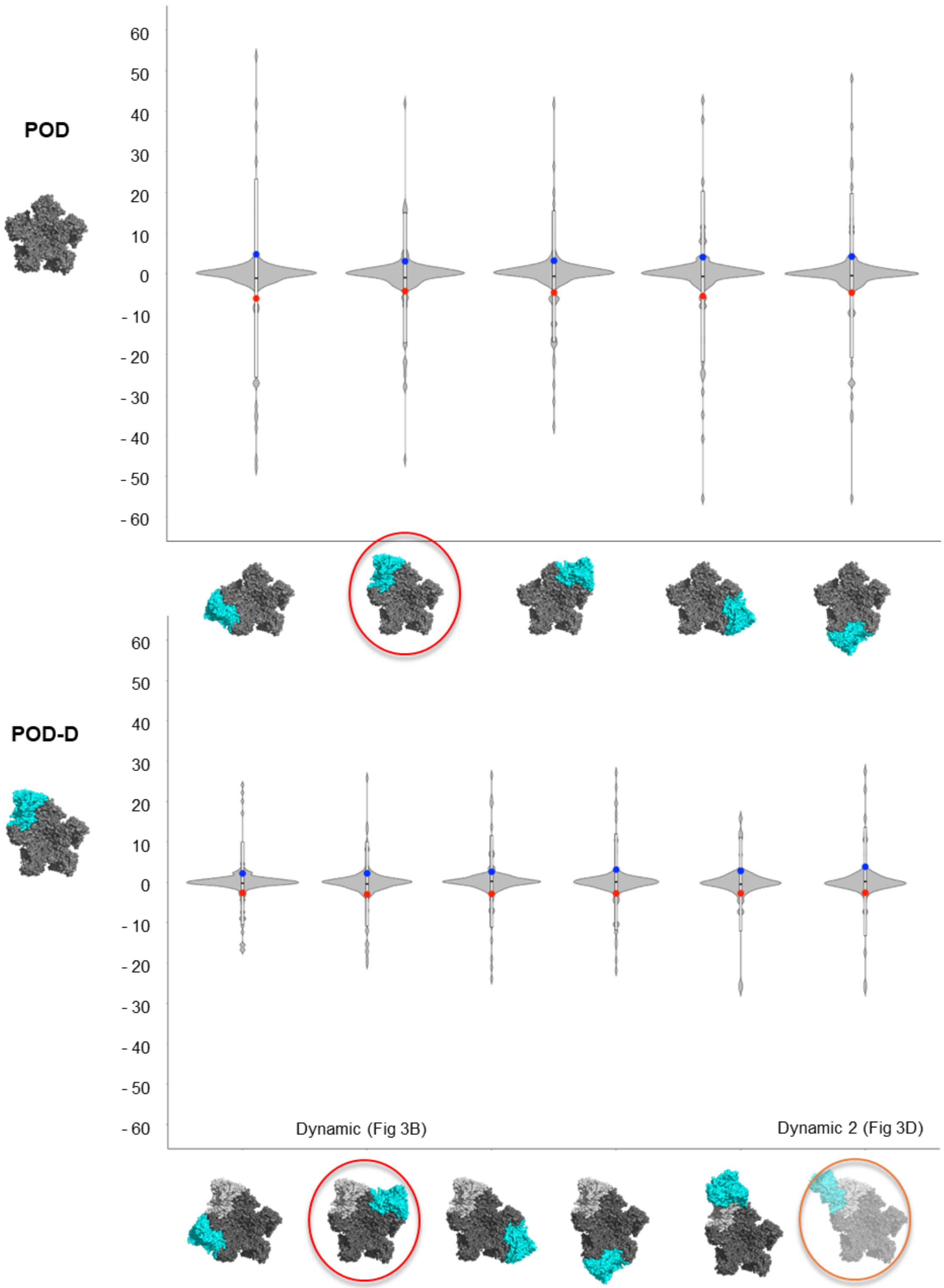


**Supplementary Figure 4.**

**Violin plots of ionizable shifted residues at the POD or POD-D docking interface.** The violin plot displays the distribution of the shifted charges of the ionizable residues. It contains a boxplot that indicates the mean in the middle and the standard deviation at the extremities. The blue point corresponds to the mean of positive shifted charges and the red point corresponds to the mean of negative shifted charges. Docking positions selected to define capsid growth are circled in red.

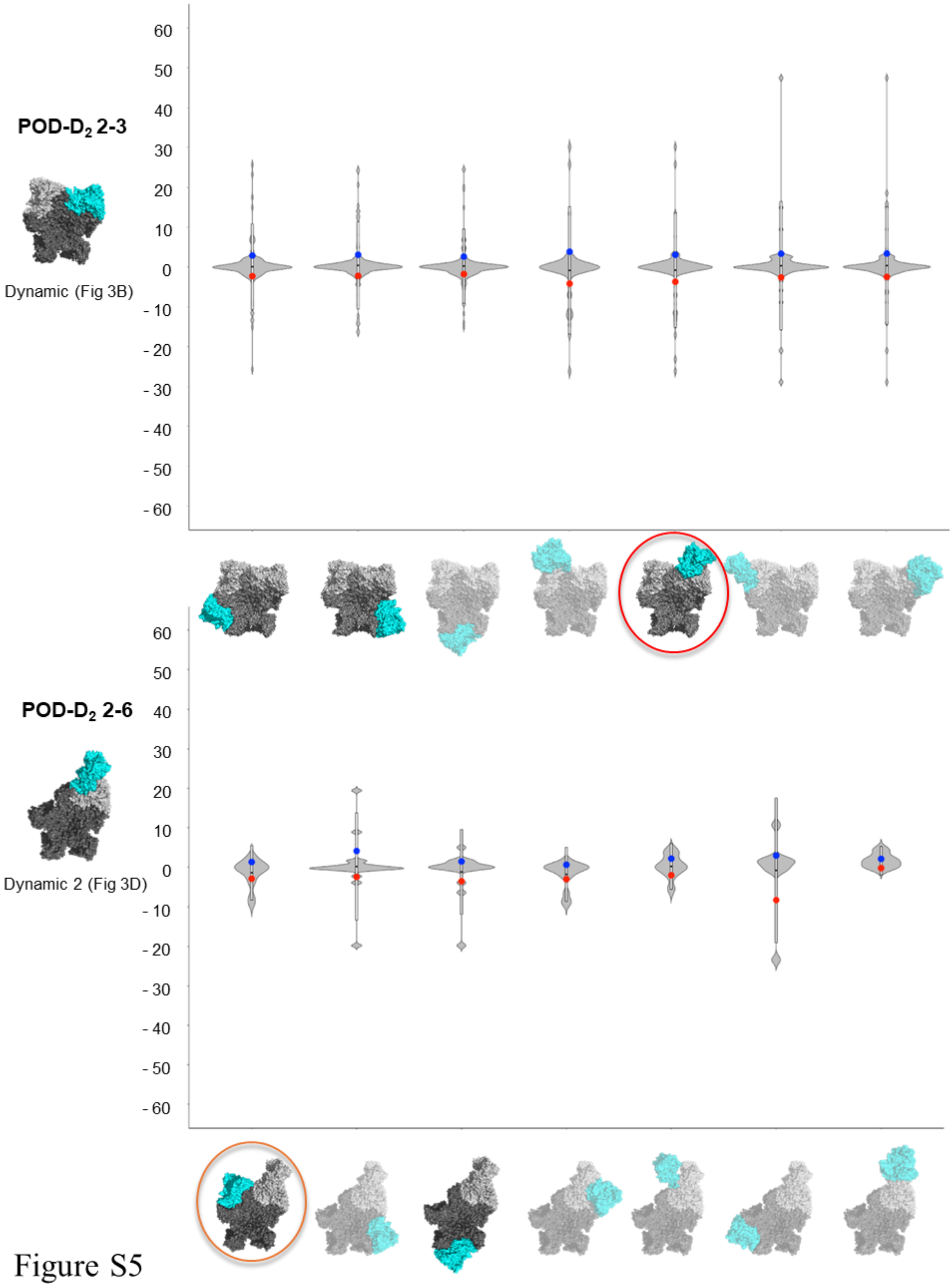


**Supplementary Figure 5.**

**Violin plots of ionizable shifted residues at the POD-$D_2$ docking interface.** The violin plot displays the distribution of the shifted charges of the ionizable residues. It contains a boxplot that indicates the mean in the middle and the standard deviation at the extremities. The blue point corresponds to the mean of positive shifted charges and the red point corresponds to the mean of negative shifted charges. Docking positions selected to define capsid growth are circled in red.

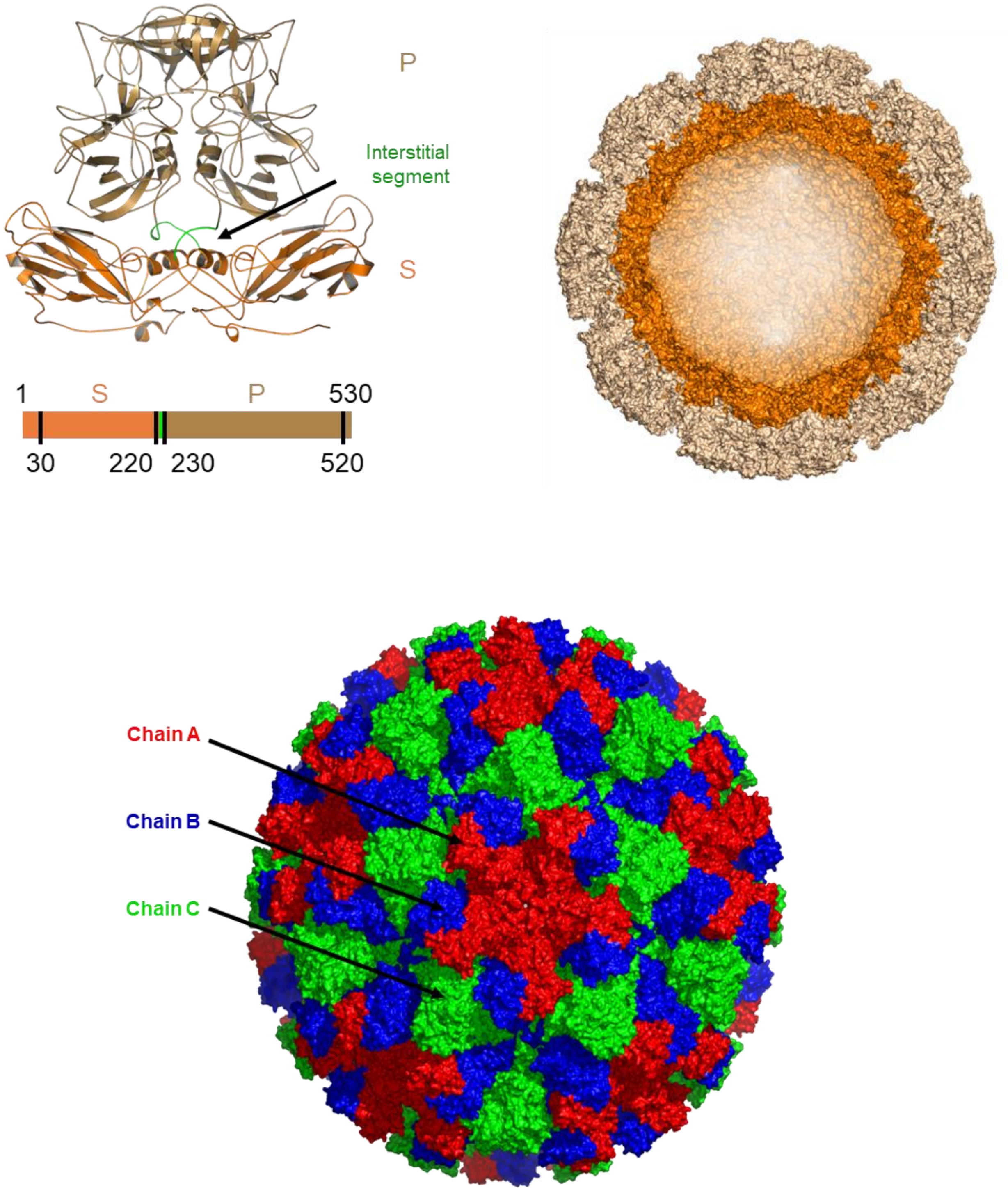


**Supplementary Figure 6.**

**Structure of VP1 dimer.** Domains S (shell, residues 20 to 220) consider as assembly domains are colored in orange, and domains P (protruding, residues 230 to 520) domain are colored in brown. The domains are connected by an interstitial loop in green. Surface and cut-away views of the capsid are displayed.
